# Characterizing Human Mesenchymal Stromal Cells Immune Modulatory Potency Using Targeted Lipidomic Profiling of Sphingolipids

**DOI:** 10.1101/2021.06.01.446428

**Authors:** S’Dravious A. DeVeaux, Molly E. Ogle, Sofiya Vyshnya, Nathan F. Chiappa, Bobby Leitmann, Ryan Rudy, Abigail Day, Luke J. Mortensen, Joanne Kurtzberg, Krishnendu Roy, Edward A. Botchwey

## Abstract

Cell therapies are expected to increase over the next decade due to increasing demand for clinical applications. Mesenchymal stromal cells (MSCs) have been explored to treat a number of diseases, with some successes in early clinical trials. Despite early successes, poor MSC characterization results in lessened therapeutic capacity once *in vivo*. Here, we characterized bone–marrow (BM), adipose derived and umbilical cord tissue MSCs’ sphingolipids (SLs), a class of bioactive lipids, using liquid chromatography – tandem mass spectrometry. We found ceramide levels differed based upon donor’s sex in BM-MSCs. We detected fatty acyl chain variants in MSCs from all 3 sources. Linear discriminant analysis revealed that MSCs separated based on tissue source. Principal component analysis showed IFN-γ primed and unstimulated MSCs separated according to their SL signature. Lastly, we detected higher ceramide levels in low IDO MSCs, indicating sphingomeylinase or ceramidase enzymatic activity may be involved in their immune potency.

## INTRODUCTION

Cellular therapies have begun to revolutionize the medical field by leading to the development of cutting-edge therapeutics to treat various diseases such as cancer, while offering promising treatments to repair damaged or destroyed tissue. As a result, the cell therapeutic industry is expected to expand over the next few decades. In 2019, the global cell therapy market was valued at $755.4 million USD and is projected to be worth $11 billion USD by 2029 ^1-4^. With the expected increase of cell therapies, scaling-up processes and manufacturing strategies have been expanded to meet clinical demand.

In particular, the manufacturing of MSCs has gained much interest due to MSCs’ multipotent differentiation, immunomodulation, immune suppression, and pro-regenerative properties via secreted cytokines, extracellular vesicles (EVs) and trophic factors ^5-8^. In recent years, preclinical and clinical trials have begun to investigate potential therapeutic effects of MSCs *in vivo*. Early studies have demonstrated that MSC-based immunotherapies show efficacy for treatment of diabetes, cardiovascular diseases, graft-versus-host disease, Crohn’s disease, and various inflammatory diseases ^9-11^. Despite pre-clinical and clinical successes, factors like culture conditions, donor history, donor variability, tissue source, manufacturing processes and cell isolation techniques affect MSC therapeutic efficacy *in vivo* ^12-14^. Minimal criteria for characterization of MSCs include adherence to plastic, expression of specific surface antigens, and multipotent differentiation potential *in vitro* ^15^, however the International Society for Cell & Gene Therapy (ISCT) has more recently called for improvements to MSC characterization by recommending the inclusion of functional assays and multi-omic profiling for additional means of characterization ^16^.

Metabolomics is an emerging field that assesses multiparametric metabolic responses of cells and tissue to external stimuli ^17^. Moreover, high throughput analytical techniques have greatly accelerated the field, allowing the collection of large and robust metabolomic datasets ^18-20^. In particular, metabolomics has been used to explore manufacturing process parameters for culture condition optimization, while offering potential advantages such as culture standardization and cost-effectiveness ^21-23^. Lipidomics, a sub-branch of metabolomics, is a powerful technique that combines high-throughput analytical methods, such as mass spectrometry (MS), and informatics to quantify and characterize cells’ lipid profile (or lipidome). Lipid components make up approximately one-third of all metabolites; thus, understanding their dynamic roles in biochemical signaling will allow researchers to prevent, diagnose and treat a wide range of human diseases ^24^. With advances of MS technologies, researchers are able to investigate thousands of lipid species that may be involved in normal cell behavior and disease pathologies. This highlights the need for investigating lipid metabolism to help develop a clear understanding of their complex functions, not only in clinical settings but also in cell manufacturing.

Bioactive lipids are a class of lipids that participate in complex cellular and molecular processes that regulate cell homeostasis, cell fate, inflammation, and many other biological processes ^25^. Functionally, bioactive lipids respond to stimuli, allowing them to regulate downstream targets of interest. As a result, bioactive lipids are involved in regulatory cell circuits, which sets them apart from other lipid classes ^26^. Sphingolipids (SLs) are a class of bioactive lipids that are comprised of a 18-20 carbon sphingosine backbone, a polar headgroup and a fatty acyl chain, offering unique functional roles in cell biology. SLs participate in various signaling pathways related to cell fate, proliferation, extracellular vesicle biogenesis, migration, differentiation, inflammation and other important processes ^27, 28^. Limited studies suggest that SLs sphingosine-1-phosphate (So1P) and ceramide-1-phosphate influence MSC’s osteogenic differentiation, morphology and proliferation ^29-31^, but no studies have measured the broader network of bioactive SL metabolites in MSCs. In this study, we characterized MSCs’ sphingolipidome using liquid chromatography – tandem MS (LC-MS/MS). We obtained MSCs derived from bone marrow (BM), adipose (AD), and umbilical cord tissue (UCT) from male and female donors and extracted their sphingolipids, as described in our previous work ^32^. This study investigated how factors like donor’s sex, culture conditions, cytokine priming, and MSC tissue origin affect the MSC SL profile. SL characterization of sex-specific differences and MSC tissue origin have not been reported. These factors may play a role in MSC *in vivo* therapeutic efficacy and should be explored for critical quality attribute optimization in the cell manufacturing sector.

## RESULTS

### LC-MS/MS Detection of LCB and CSLs in MSCs

Sphingolipids (SLs) are a structurally diverse class of membrane lipids that are composed of an 18 to 20 carbon amino-alcohol backbone, sphingosine (So), and are synthesized in the endoplasmic reticulum (ER) ^33^. So and dihydrosphingosine, also referred to as sphinganine (Sa), are the basic building blocks of SLs in mammalian cells. This “sphingoid” backbone can be modified to produce a wide variety of SLs with distinct functions in cell signaling and structural adaptations of biological membranes (**Figure 1, Table 1**) ^34^. N-acylation of sphingosine by ceramide synthases (CerS) generates ceramide (Cer), whereas phosphorylation of the C1-hydroxyl group of sphingosine by sphingosine kinase (SK) produces sphingosine-1-phosphate (So1P) ^35^. Cer can in turn serve as a precursor of more complex sphingolipids (CSLs), such sphingomyelins (SMs) and ceramide 1-phosphate.

**Table 1.**
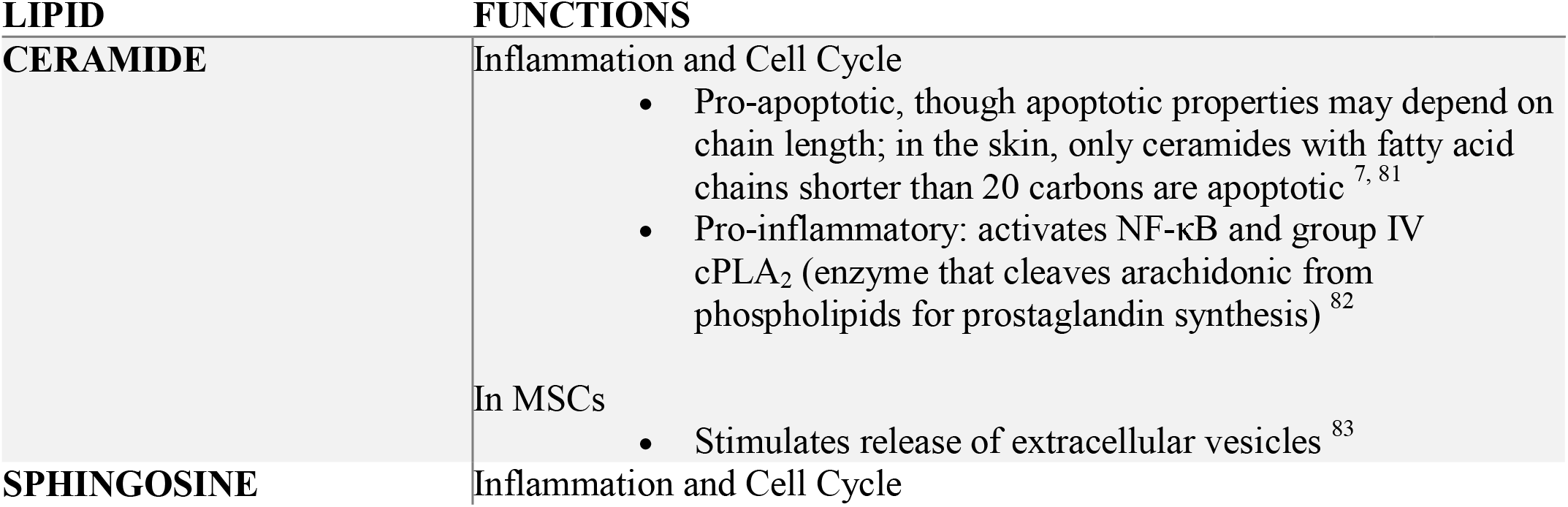
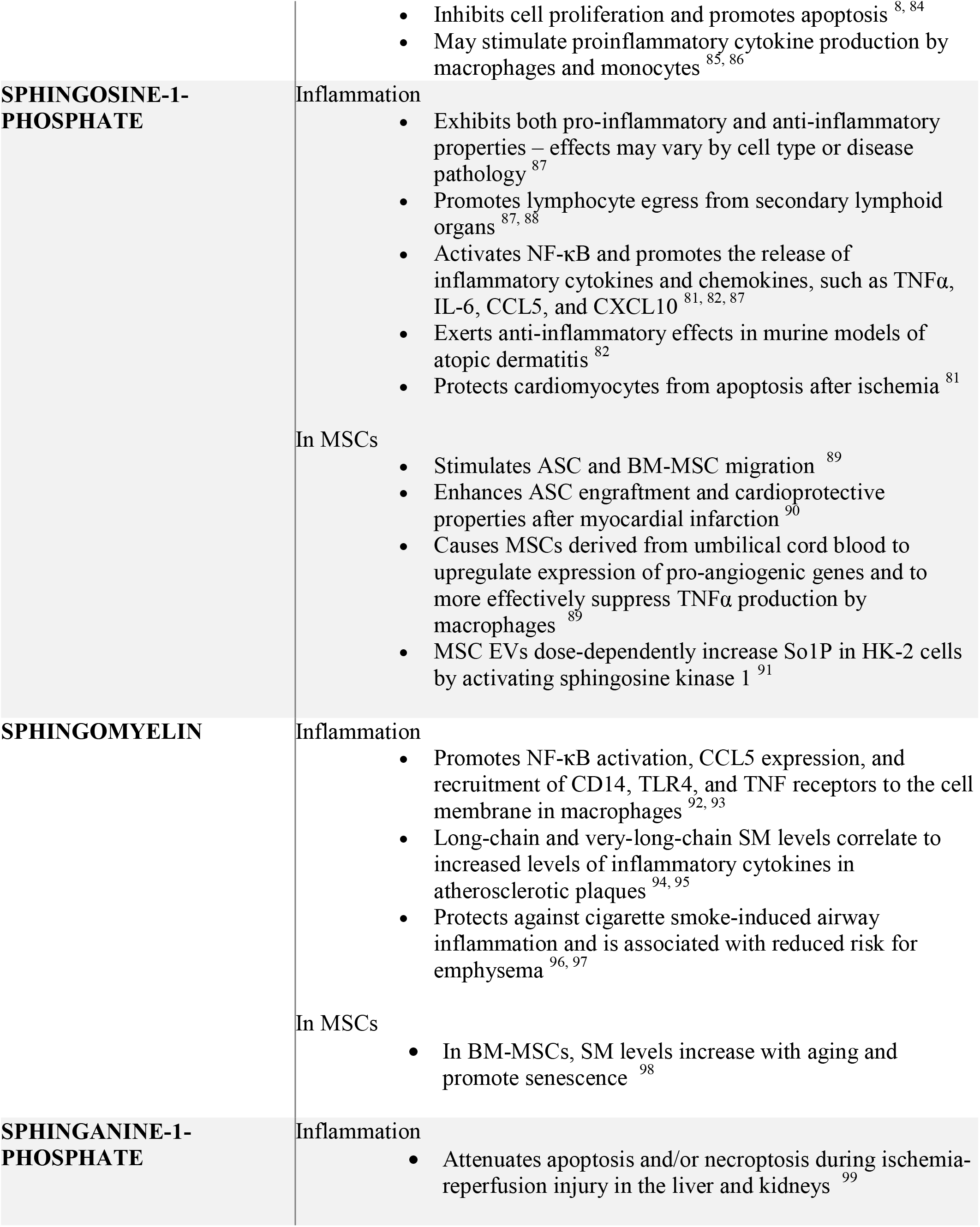
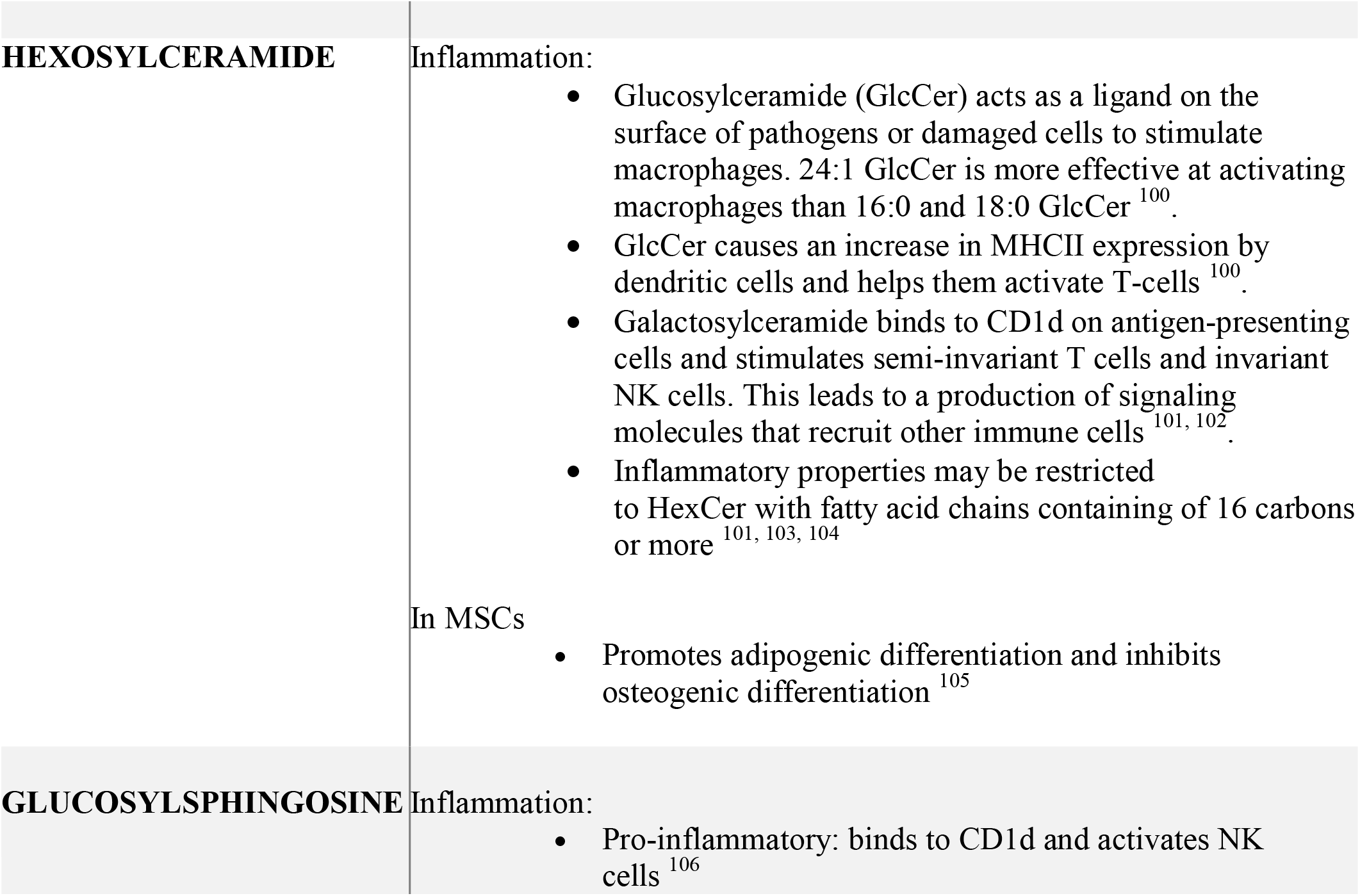
Sphingolipid functions.

**Figure 1.**
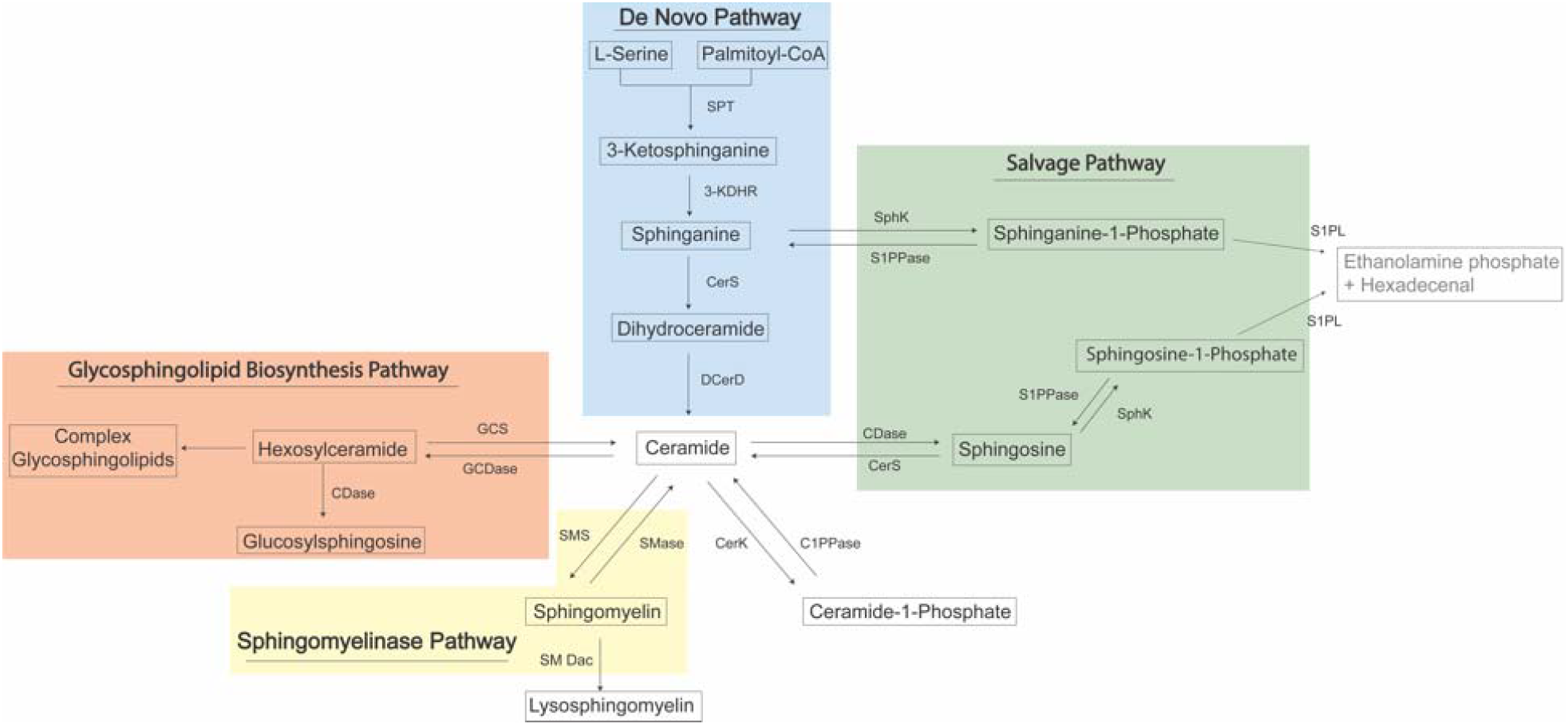
Sphingolipid metabolic pathway. SPT Serine Palmitoyltransferase; 3-KDHR 3-Ketosphinganine Reductase; CerS Ceramide Synthase; DES Dihydroceramide Desaturase; CDase Ceramidase; GCS Glucosyl-/Galatosyl-ceramidase; SphK Sphingosine Kinase; S1PPase Sphingosine-1-Phosphate Phosphatase; SMS Sphingomyelin Synthase; SMase Sphingomyelinase; SMDac Sphingomyelin Deacylase; CerK Ceramide Kinase; C1PPase Ceramide-1-Phosphate Phosphatase.

To evaluate the MSC’s sphingolipidome, we obtained MSCs derived from bone-marrow (BM), adipose (AD), and umbilical cord tissue (UCT), conducted SL extraction, and measured the lipid amounts with LC-MS/MS in multiple reaction mode. Precursor and product ions were input prior to MS analysis for targeted analysis. Long chain bases (LCBs) and CSLs amounts were normalized by total protein to facilitate MSC tissue source comparison. We detected and measured the following LCBs: Sa, Sphinganine-1-Phosphate (Sa1P), So, Glucosylsphingosine (GlcSo), So1P and Lysosphingomyelin (LSM). We detected and measured the following CSLs: SM, Cer and Hexosylceramide (HexCer) (**Figure 2**).

**Figure 2.**
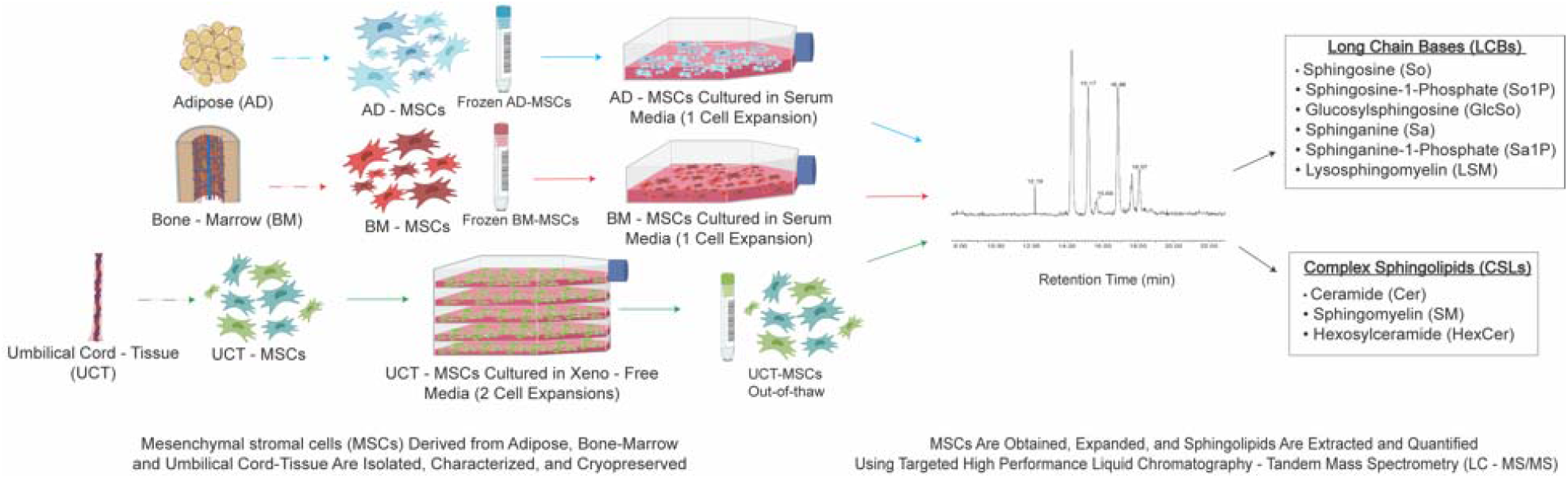
Experimental overview of study

When assessing UCT-MSCs’ SL profile, it consisted of 99.75% CSLs and 0.25% LCBs. The CSL pool was comprised of Cer (38.65%), SM (35.19%), then HexCer (25.91%). When measuring the LCBs, Sa1P was not detected in UCT-MSCs. The So and So1P made up majority of the LCB pool with 47.35% and 31.10%, respectively. Sa (18.75%), GlcSo (2.38%) and LSM (0.43%) made up the remaining detected LCBs (**Figure 3A**). AD-MSCs’ total sphingolipidome consisted of 98.04% CSLs and 1.60% LCBs. SM was the predominant CSL detected (73.45%), followed by Cer (22.89%) and HexCer (2.05%). Six LCB species were detected in AD-MSCs. So (64.59%) was the predominant LCB detected. The least detected LCB, LSM only made up 0.21% of the LCB lipid pool. The remaining LCB pool was made of Sa (22.20%), Sa1P (7.69%), GlcSo (0.70%), and So1P (4.61%) (**Figure 3B**). Comparing the SL distribution between sexes, male and female BM-MSCs had observable differences. Male BM-MSCs’ sphingolipidome consisted of 54% Cer, 44.53% SM, 1.48% HexCer and 1.48% LCBs. The LCBs consisted of 23.64% So, 8.04% So1P, 28.07% Sa, 6.18% Sa1P, 21.84% GlcSo and 12.22% LSM (**Figure 3C**). Contrastingly, the female BM-MSC sphingolipidome was comprised of 88. 35% Cer, 10.18% SM, 0.68% HexCer and 0.61% LCBs. The LCB pool was made up of 16.69% Sa, 2.57% Sa1P, 76.00% So, and 0.15% LSM (**Figure 3D**).

**Figure 3.**
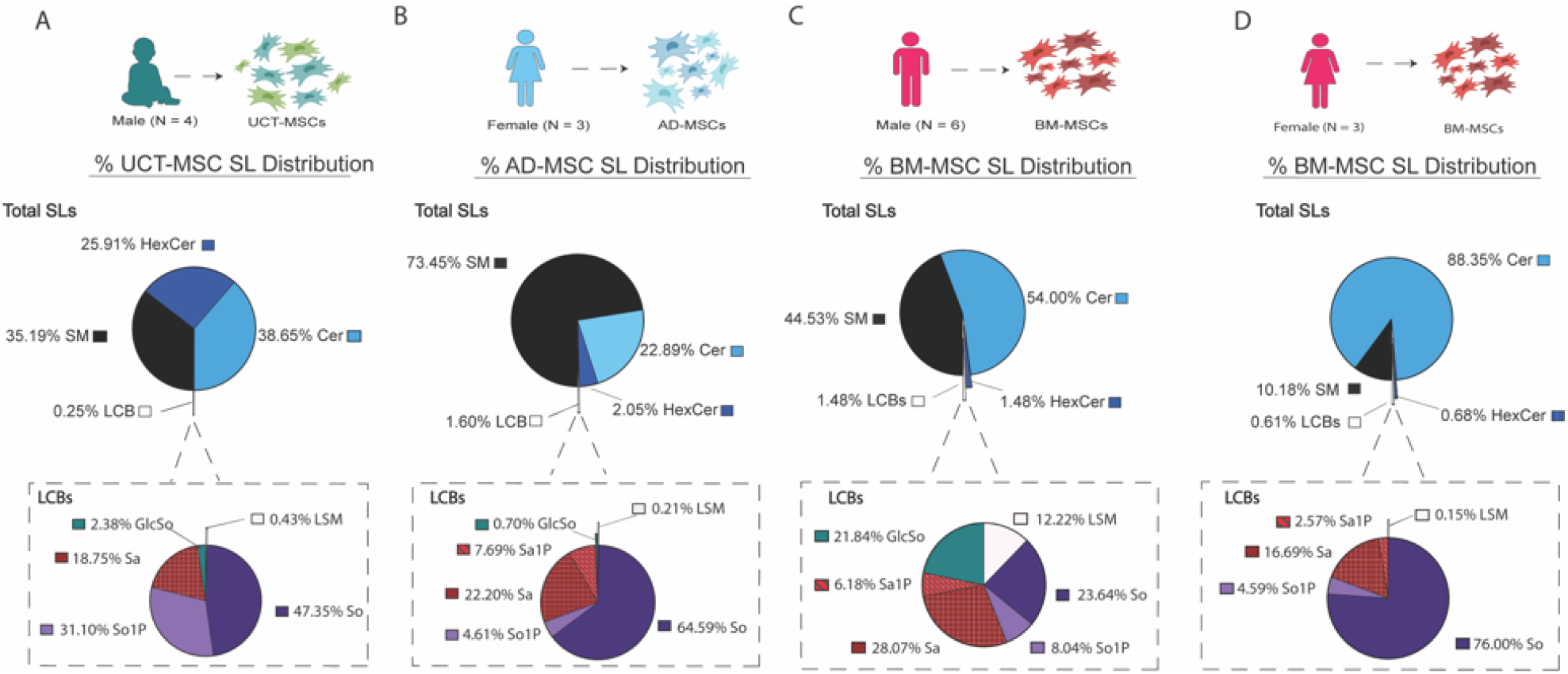
LC-MS/MS detects CSLs and LCBs in MSCs from all tissue origins. Pie chart illustrates the total SL distribution and the call-out pie charts illustrates the LCBs. A) Percent SL species distribution in UCT-MSC (N = 4). B) Percent SL species distribution of AD-MSCs (N = 3). C) Percent SL species distribution of male BM-MSC donors (N = 6). D) Percent SL species distribution of female BM-MSC donors.

### SPHINGOLIPID ACYL CHAIN VARIANTS

More recently, it has been shown that the functions of SLs differ due to the length of their fatty acyl chains ^36, 37^. Fatty acyl chain variants have not been well characterized in MSCs derived from different tissues. To assess the various SL fatty acyl chain variants, SLs were extracted and quantified with LC-MS/MS. Fatty acyl chain lengths ranging from 16-26 carbon chains containing 0-2 double bonds were detected in MSCs from all tissue sources. **Figure 4A** illustrates SM acyl chain variants detected in AD-, BM- and UCT-MSCs. SM fatty acyl chain lengths 16:0, 20:0, 22:0, 24:0 and 26:0 were not detected in AD-MSCs.

**Figure 4.**
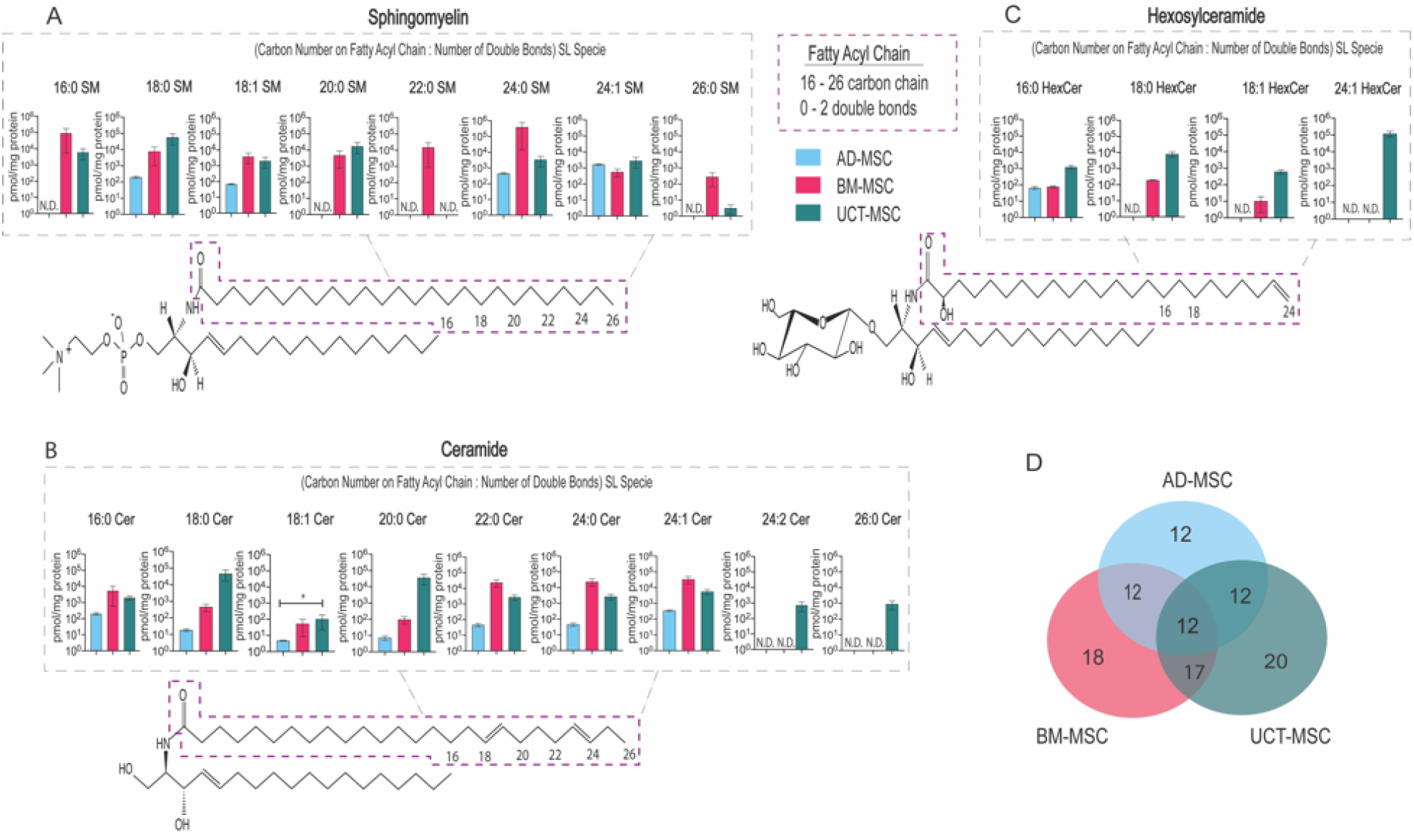
Fatty acyl chain lengths variants detected by LC-MS/MS in AD-, BM- and UCT-MSCs. Every sphingolipid consists of an 18-20 carbon sphingosine backbone, a polar headgroup and a fatty acyl chain. The purple box highlights the fatty acyl chain location on SM, Cer and HexCer. A) SM fatty acyl chain variants. B) Cer fatty acyl chain variants. C) HexCer fatty acyl chain variants. D) Venn-diagram of the number of fatty acyl chain lengths variants detected in the 3 tissue origins. 21 fatty acyl chain variants were measured. The overlap between tissue origins are the shared fatty acyl variants; N.D. not detected; AD-MSC (N = 3); BM-MSC (N = 9); UCT-MSC (N = 4). An asterisk (*) denotes statistical significance (p < 0.05). Kruskal-Wallis test and Multiple comparison were used to determine significance.

BM-MSCs had greatest amounts of 16:0 SM and 22:0 SM, while 18:0 SM was primarily detected in UCT-MSCs. BM-MSCs and UCT-MSCs had similar 18:1 SM and 20:0 SM concentrations. All three MSC groups had comparable concentrations of 24:1 SM. 24:2 SM was not detected in AD-, BM-, or UCT-MSCs (data not shown). **Figure 4B** shows Cer acyl chain lengths detected in AD-, BM- and UCT-MSCs. UCT-MSCs had the highest concentration of 18:0 Cer, 20:0 Cer, 24:1 Cer, and 26:0 Cer compared to AD- and BM-MSCs. UCT-MSCs had statistically significantly more 18:1 Cer than AD-MSCs. BM-MSCs had slightly higher concentrations of 16:0, 22:0, 24:0 and 24:2 Cer compared to AD-MSCs and UCT-MSCs. Fatty acyl chains 24:1 Cer and 26:0 Cer were not detected in AD- and BM-MSCs. **Figure 4C** shows HexCer acyl chain lengths detected in AD-, BM- and UCT-MSCs. UCT-MSC had the highest concentrations of 16:0 HexCer, 18:0 HexCer, 18:1 HexCer and 24:1 HexCer. AD- and BM-MSCs had similar levels of 16:0 HexCer. 18:0 HexCer, 18:1 HexCer and 24:1 HexCer were not detected in AD-MSCs. Additionally, 24:1 HexCer was not detected in BM-MSCs. **Figure 4D** shows the number of SL acyl chain length variants detected in AD-, BM- and UCT-MSCs. There was a total of 21 acyl chain metabolites detected and measured by LC-MS/MS, illustrated in Figures 4A-C. 12 out of 21 metabolites were detected and measured in AD-MSCs. These 12 metabolites were also measured in both BM-MSC and UCT-MSCs. 18 out of 21 fatty acyl chain variants were measured in BM-MSCs. 20 out of 21 fatty acyl chain variants were measured in UCT-MSCs. There were 17 common fatty acyl chain variants measured in both BM- and UCT-MSCs. 12 fatty acyl chain variants were measured in MSCs from all 3 tissue sources.

### ROBUSTNESS OF SL CONCENTRATIONS ACROSS TISSUE SOURCES

To understand the relationship between SL concentration and MSCs tissue source, Spearman correlation was conducted. The MSC tissue source (the independent variable) and the SL concentration (the dependent variable) were used to generate heatmaps in Python Seaborn. A monotonic relationship illustrating a strong negative association was illustrated by a linear, downward slope. Interestingly, AD-MSC donors were strongly correlated with one another, with donors AD2 and AD3 being more strongly correlated than AD1. The BM-MSCs were all strongly correlated with one another based upon their SL signature illustrated by warmer colors. Notably, a strong correlation was evident between the BM- and AD-MSCs. This could be a result of another of cell type similarity or another underlying factor, such as culture conditions.

Uniquely, the UCT-MSC donors (with the exception of UCT1a and UCT1b) were not strongly correlated with one another, as evidenced by cooler colors between the donors. Donor UCT1a and UCT1b came from the same donor but were processed at different times. There was a strong correlation between these donors, as expected (correlation coefficient ≥ 0.95) (**Figure 5A**). When comparing MSCs from the same tissue source, the MSCs were all strongly correlated with one another, illustrated by warmer colors and high correlation coefficient values (correlation coefficient ≥ 0.95) (**Figure 5B-D**). To gain insight on whether the MSCs can be classified into their tissue origin based upon their SL profiles, LDA was performed. Clear separation between the donors three tissue sources occurred. It can be inferred that donor similarities in the SL compositions led to class separation based on tissue source (**Figure 5E**).

**Figure 5.**
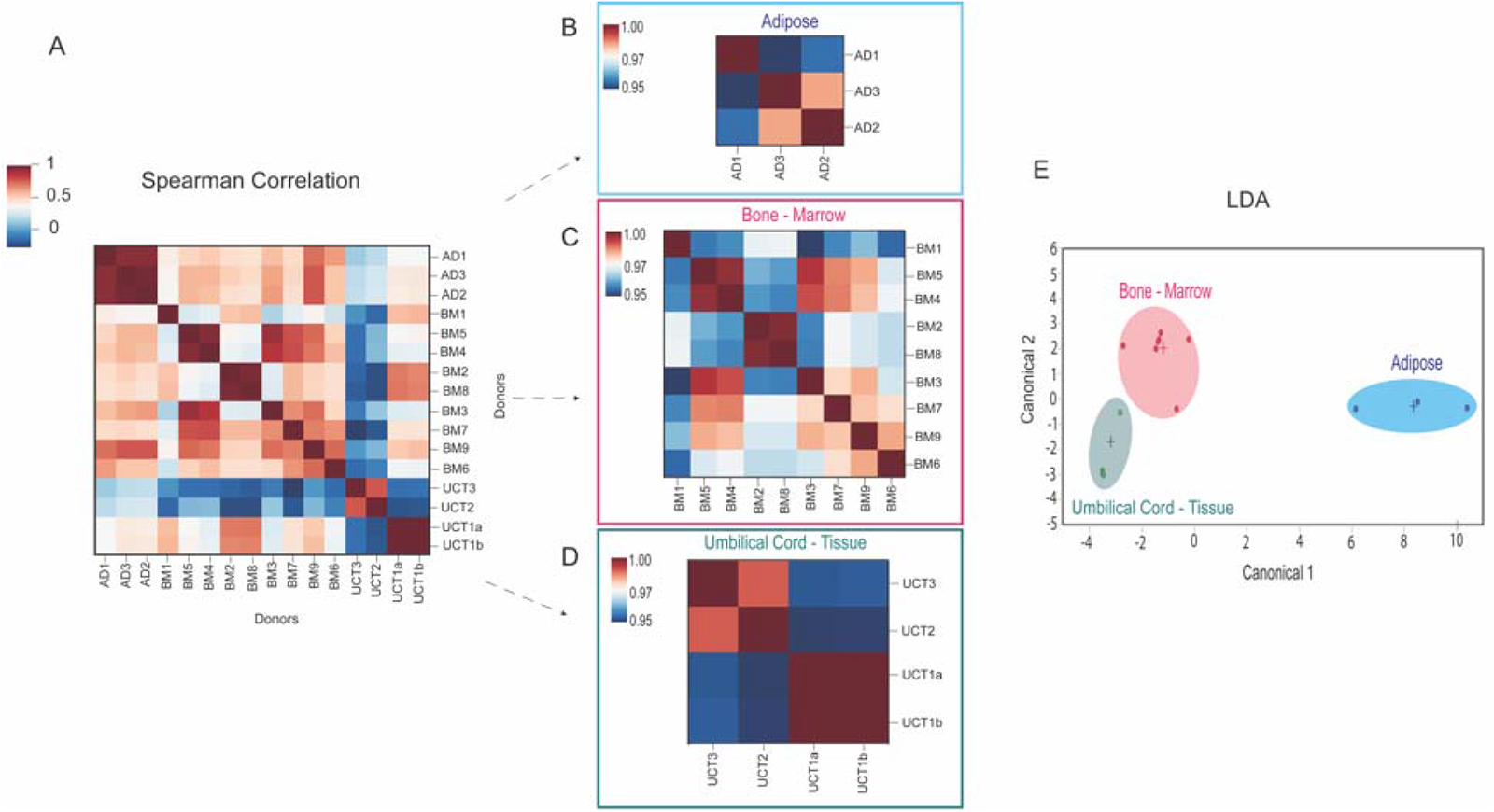
Heatmap spearman correlations and LDA of MSC tissue sources. A) Spearman’s correlation heatmap of MSC donors from the 3 tissue origins B) Spearman’s correlation heatmap of AD-MSC donors C) Spearman’s correlation heatmap of BM-MSC donors D) Spearman’s correlation heatmap of UCT-MSC donors E) LDA of MSCs donors from the 3 tissue origins. AD-MSC (N = 3); BM-MSC (N = 9); UCT-MSC (N = 4).

### DIFFERENCES IN HIGH AND LOW IDO BM-MSCS

Indoleamine 2,3-dioxygenase (IDO) activity is a common benchmark of immunomodulatory potency. This enzyme catalyzes the conversion of tryptophan to kynurenine, which is an immunosuppressive metabolite, inhibiting pro-inflammatory immune cell activity ^38^. Hence, high IDO activity (> 30 pg kynurenine/cell/day) indicates higher immunomodulatory potency (**Figure S1**). We stimulated MSCs from two BM donors (BM1 and BM6) with IFN-γ, which resulted in an increase of IDO enzymatic activity ^39^. Based on the IDO activity values provided by the manufacturer, donor BM1 was classified as low-IDO, as its IDO activity was below the mean (30 pg kynurenine/cell/day), and donor BM6 was classified as high-IDO, as its IDO activity was above the mean (**Figure S1**) (**Figure 6A**).

**Figure 6.**
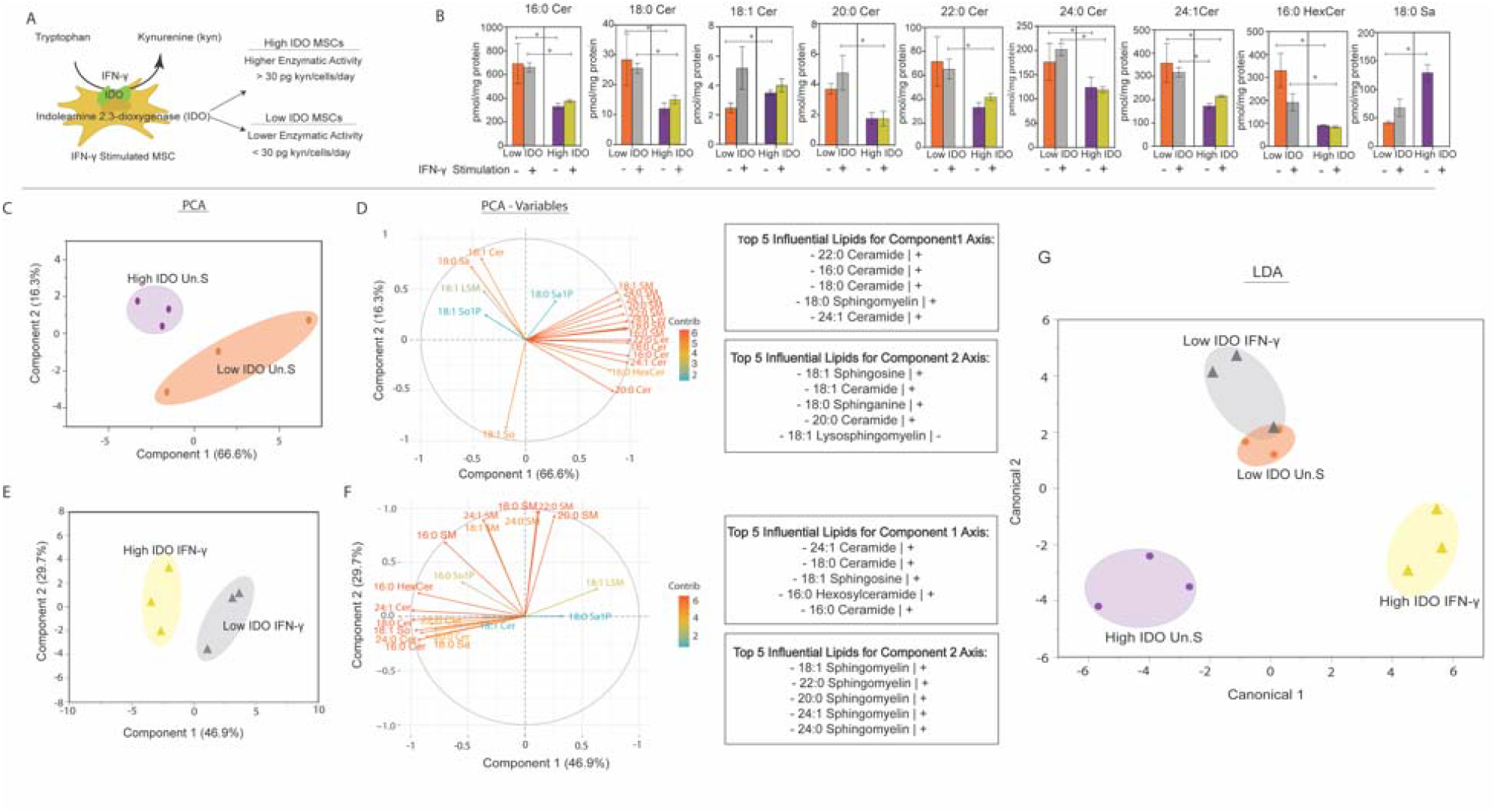
Unstimulated and IFN-γ stimulated BM-MSCs SL profile. A) Schematic illustrating indoleamine 2,3-dioxgenase (IDO) converting tryptophan to kynurenine (kyn), an immunosuppressive metabolite. B) SL concentrations of unstimulated and IFN-γ primed MSCs C) PCA of high and low IDO unstimulated (Un.S) MSC groups D) PCA variable plot of MSC groups. E) PCA of high and low IDO IFN-γ primed groups. F) PCA variable plot of IFN-γ groups. G) LDA of high and low MSCs from Un.S and IFN-γ group-s. An asterisk (*) denotes statistical significance (p < 0.05). Mann-Whitney U-test was used to determine significance. All groups were measured in triplicates (n = 3).

We investigated the relationship between IDO activity levels and the MSC sphingolipidome by comparing LC-MS/MS analysis profiles of unstimulated (Un.S) and IFN-γ primed high and low IDO BM-MSCs. Principal component analysis (PCA) was conducted to determine whether BM-MSCs can separate into high- and low-IDO groups based on their SL profiles. (**Figure S1**). When measuring SL species, greater concentrations of acyl chain Cer species (16:0, 18:0, 20:0, 22:0, 24:0 and 24:1) and 16:0 HexCer were detected in both low IDO MSC treatment groups compared to both high IDO MSC treatment groups. Unstimulated high IDO had the greatest concentration of 18:0 Sa; however, Sa was not detected in high IDO IFN-γ treated MSCs. When looking at the effect of priming MSCs with IFN-γ on the SL profile, low IDO MSCs had greater concentrations of 18:1-, 20:0-, 24:0 Cer and 18:0 Sa compared to the low IDO unstimulated group. The remaining SL species had comparable levels, with the exception of 16:0 HexCer, where greater amounts of SLs were detected in the unstimulated group. In high IDO MSCs, unstimulated and IFN-γ primed MSCs had comparable concentrations of SLs, with the exception of Sa (**Figure 6B**). IFN-γ priming did not significantly alter SM levels in high or low IDO groups.

To understand the relationship between SL species detected in high and low IDO BM-MSCs, PCA was conducted. The PCA score plot of unstimulated MSCs shows a clear separation between high-IDO and low-IDO groups. The separation was primarily due to the first principal component (PC1), which explains 66% of the variance in the data, and the second principal component (PC2), which explains 16% of the variance (**Figure 6C**). The lipids that most strongly contributed to the separation along PC1 predominantly consist of ceramide species (22:0 Cer, 16:0 Cer, 18:0 Cer, and 24:1 Cer), as well as one sphingomyelin species (18:0 SM) (**Figure 6D**). The lipids with the greatest influence on separation along PC2 include 18:1 Cer,18:0 Sa, 18:1 So, 20:0 Cer, and 18:1 LSM. All the lipids except for 18:1 LSM correlate positively along the PC2 axis. These results suggest possible differences in ceramide and LCB metabolism between high IDO and low IDO MSCs in the unstimulated state. When conducting PCA on IFN-γ primed high and low IDO groups, separation was primarily due to PC1, with 46.9% of the variance in the data, and PC2 with 29.7% (**Figure 6E**). The lipids that most strongly contributed to the separation along PC1 consisted of 24:1 Cer, 18:0 Cer, 18:1 So, 16:0 HexCer, and 16:0 Cer. The lipids with the greatest influence on separation along PC2 were predominantly SM species (18:1 SM, 22:0 SM, 20:0 SM, 24:1 SM and 24:0 SM) (**Figure 6F**). This suggests differences in ceramide and sphingomyelin metabolism between low IDO and high IDO MSCs after IFN-γ priming.

Furthermore, we wanted to assess whether SLs can be used as predictor features to classify MSCs into groups based on their IDO activity. We utilized linear discriminant analysis (LDA), which is a supervised machine learning algorithm that can perform dimensionality reduction and classification. When used for dimensionality reduction, LDA relies on the class labels of each observation and projects the data onto an axis that maximizes the separation between the classes ^40^. LDA is increasingly being used in biomedical applications to differentiate between healthy and diseased states or to classify various cell states within the same cell line ^41-43^. We utilized SL concentrations as the input features and the IDO activity + IFN-γ treatment (low IDO unstimulated, low IDO IFN-γ primed, high IDO unstimulated, high IDO IFN-γ primed) as the class labels in LDA. The LDA plot showed separation between high IDO unstimulated and IFN-γ groups. Interestingly, the low IDO unstimulated and low IDO IFN-γ did not have much separation, possibly due to similar SL profiles (**Figure 6G**).

## DISCUSSION

MSCs have shown to be therapeutic in preclinical and early clinical trials in treating acute and chronic inflammatory diseases. Despite their therapeutic potential, discrepancies MSC characterization have limited further clinical application. Our research is motivated by a growing appreciation of how local accumulation of specific sphingolipids (SLs) either directly or indirectly regulates multiple cellular properties that may predict stem cell potency, including mechanics of plasma and nuclear membrane, mitochondrial activity, and cell death mechanisms ^35, 44, 45^. SLs are a highly diverse family of molecules which are important building blocks of eukaryotic membranes ^33^. Prior to this work, we demonstrated that modulating novel sphingosine-1-phosphate (S1P) and S1P receptor 3 (S1PR3) signaling mobilizes both hematopoietic stem cells (HSCs) and MSCs to enhance ectopic bone growth, further emphasizing the importance of understanding SL functions ^46, 47^. In this study, we performed LC-MS/MS to characterize AD-, BM-, and UCT-MSCs sphingolipid metabolism profiles.

Demographic features have not been fully explored as it pertains to MSC lipidomic characterization. It is not well understood how factors like donor’s sex affect MSC SL metabolism. When comparing male and female MSC ceramide (Cer) levels, females possess greater levels of Cer. Generally, Cer, a central intermediate, regulates cell fate by activating the apoptosis regulator proteins, Bcl-2-associated X and Bcl-2 homologous antagonist, leading mitochondrial outer membrane permeabilization and apoptosis ^48^. This internal feedback system illustrates SL complex interconnectivity and signaling. It is also interesting to speculate that differences in sphingolipid profiles are a result of differences in body composition of male and female donors and/or differing hormonal influences on cell properties. ^49-51^. Li et al. recently reported relationship between Cer and estrogen, showing decreased Cer synthase (CerS) and sphingomyelinase activity in ovariectomized rats. This resulted in perimenopausal hypertension, suggesting possible changes Cer homeostasis among female donor cells ^52^. As we detect SL differences in BM-MSCs from male and female donors, additional studies and analysis investigate the effect of sex on MSC sphingolipidome are needed.

Priming MSCs with IFN-γ activates their anti-inflammatory functions, thus priming them for immunomodulation ^53^. IFN-γ increases MSCs indoleamine 2,3-dioxygenase (IDO) enzymatic activity, which catalyzes the rate-limiting step in the conversion of the amino acid tryptophan into kynurenine ^54, 55^. MSCs with higher IDO activity (≥ 30 pg kyn/cell/day) are more effective in regulating immune responses than MSCs with lower IDO activity (< 30 pg kyn/cell/day) ^56^. However, variability in immune modulatory potency among donors based on IDO assays remains a clinical hurdle ^57^. When investigating the effects of interferon-gamma (IFN-γ) on MSCs, Campos et al. uncovered increases in very-long chain (34-42 carbon chain) sphingomyelin (SM) species after priming ^31^. However, these studies did not solely focus on MSC sphingolipidome characterization. Assessing the sphingolipid profile of unstimulated and IFN-γ primed BM-MSCs uncovered differences in Cer species and Sa between low and high IDO MSCs. Cer acts as a secondary messenger in signaling pathways relating to immune responses such as cytokine production and cell activation ^58^. Jrad-Lamine et al. showed mice lacking the expression of *Ido1* gene possessed higher levels of Cer compared to wild type mice ^59^. Our findings show a similar trend, where low IDO MSCs have higher Cer levels. It remains unclear whether and how changes in Cer signaling may influence immunomodulatory properties or paracrine activity of MSCs directly, but these results do raise the exciting possibility that pharmacological or genetic inhibition of enzymes resulting in Cer accumulation, such as sphingomyelinase, may be effective in culture to preserve or enhance the therapeutic potency of MSCs. Further studies will need to be conducted to understand the role of Cer in low IDO MSCs and associated implications on their *in vivo* immune modulatory properties.

Circulating lipids, such as cholesterols and triglycerides, have been the gold standard for understanding the progression of diseases, such as hypertension, diabetes and cardiovascular diseases ^60, 61^. However, SLs in particular have a distinct role where they form cellular membranes together with glycerolipids and sterols. This allows regulation of sorting and trafficking of membrane content and regulation of cellular morphology ^62, 63^. Sphingomyelins (SMs) are the most abundant SL in the membrane of mammalian cells, where they facilitate integral protein recruitment via SM-cholesterol interactions, lipid raft formation, regulation of endocytosis, ion channel flux regulation, and modulating the biophysical properties of the cell membrane ^64, 65^. We detected SMs primarily in AD-MSCs. *In vivo*, adipocytes rely on lipid metabolism to regulate their unique functions. In particular, structural membrane SLs such as SM, Cer, and So1P regulate intracellular mechanistic roles influencing adipocyte morphology and lipid droplet biogenesis. Alexaki et al. also demonstrated that introducing perturbations to the SL de novo pathway, resulted in lower SM levels and a hypertrophic phenotype common to lipodystrophies and apoptotic adipocytes ^66^. It can be inferred from our analysis that MSCs derived from mature tissue sources such as AD-MSCs may retain characteristic SL profiles and cellular properties after isolation from adipose tissue. Previous lipidomic analyses of white adipose tissue wherein differing types of fat derived from either the visceral or subcutaneous fat sources have shown that the tissues themselves retain a distinct “fingerprint” of lipid concentration ^67^; and so, our findings imply that characterization methods targeting lipid metabolism in general and bioactive lipids in particular is a promising approach to assess therapeutic quality and potency of cell therapies.

SLs play an essential role in multiple stages during prenatal development and tissue maturation. Glycosphingolipids along with So1P participate in endocytosis of nutrients in the intestines, skin barrier permeability and homeostasis, and neural development ^68-71^. In our study, we found that glycosphingolipids like HexCer and LCB So1P had higher concentrations detected in UCT-MSCs. These distributions may be attributable to their role in developmental maturity of umbilical cord tissue, where cells derived from more primitive tissue sources retain characteristic SL metabolic profiles. Biosynthesis of individual fatty acyl chain variants of SLs species may also be predictive of tissue source and cell potency. Six (dihydro) CerS isoforms have shown fatty acyl-CoA substrate specificity leading to variations of the ceramide molecule based on saturation (saturated or unsaturated) or length (C14–C26) of the N-acyl chain. For example, CerS1 produces C18-Cer species with stearoyl CoA as a substrate, while CerS2 is responsible for producing very long chained fatty acyl Cer species (C20-26) ^72^. CerS3 prefers middle and long chain fatty acyl CoAs to generate large structural SLs ^73^ whereas CerS5 and CerS6 prefer palmitoyl-CoA substrate to generate C16 Cer species ^74, 75^. Therefore, MS-based determination of Cer acyl chain variations in cultured MSCs may elucidate which isoforms of CerS that are most active. Interestingly, CerS2 and CerS3 expression is elevated in more primitive cells, including the placenta (CerS2) and testis (CerS3), and more weakly expressed in more mature tissues ^75, 76^. These findings suggest generation of SL profiles may reveal novel enzymatic targets associated with cell maturity changes in culture. As specific acyl chain variants of Cer affect diverse biological processes ranging from embryo development, macrophage differentiation and tumor progression ^77^, it is possible that by elucidating the role of CerS isoforms on cultured MSCs, we may determine whether generation of acyl chain variants of SLs only correlate with immunomodulatory properties or influence MSC properties directly.

In conclusion, we used state of the art LC-MS/MS sphingolipidomic approaches established at Georgia Tech / Emory ^78-80^ to study how variations in the concentrations of regulatory SL metabolites may be used to characterize the quality and potency of cultured MSCs. Multi-variate analysis of lipidomic network profiles yielded valuable insight into tissue origin and immune modulatory of the cells. This analysis also elucidates particular enzymatic targets in the sphingolipid metabolic network that may be targeted to minimize undesired changes in membrane composition during culture to preserve or enhance cell quality.

## ACKNOWLEDGEMENTS

This work was funding by The Marcus Foundation, the Marcus Center for Therapeutic Cell Characterization and Manufacturing, The Georgia Tech Foundation, and the Georgia Research Alliance. Krishnendu Roy was supported by the National Science Foundation Engineering Research Center for Cell Manufacturing Technologies (NSF EEC 1648035). S’Dravious DeVeaux was supported by the NGMS-sponsored Cell and Tissue Engineering NIH Biotechnology Training Grant (T32 GM-008433). We would like to thank Research Coordinator, David Bostwick in Parker H. Petit Institute for Bioengineering and Biosciences mass spectrometry core at Georgia Tech for his assistance with mass spectrometry method development. We would like to thank Kuang-Drew Li and Sami Belhareth for preliminary lipidomic analysis. Graphics used for figure illustrations were created using Biorender.com. The graphics were exported with a paid Biorender.com subscription.

## AUTHOR CONTRIBUTIONS

S.A.D. analyzed the data, generated the figures, and wrote the paper. S.V. analyzed the data. M.E.O. designed and conducted the experiments. N.F.C. developed the mass spectrometry method, performed lipid extractions and conducted the mass spectrometry lipid analysis. R.R., and A.D. performed the cell lipid extractions. B.L. and L.J.M. assisted with experimental design. J.K. supervised primary cell isolation and characterization. K.R., and E.A.B. supervised the project.

## METHODS

### UCT-MSC Isolation and Expansion

We used umbilical cord tissue-derived MSC (UCT-MSC) donors UCT1a, UCT1b, UCT2 and UCT3 for lipidomic analysis. Donors UCT1a and UCT1b were obtain from the same donor. All the UCT-MSCs were collected from the umbilical cords of male babies and expanded up to passage 2 (P2) by the Department of Pediatrics at Duke University. For a typical expansion of UCT-MSC, cryopreserved P0 vials were thawed at 37°C in water batch for up to 2 minutes and quickly transferred to a 15-ml sterile tubes containing xeno-free serum-free media (XS-FM, Prime-XV, Irvine Scientific) and gently mixed with the media by pipetting up and down. Cell viabilities and counts were measured using acridine orange-propidium iodide (AOPI) method using a NucleoCounter (Chemometec). Then, the cells were suspended in XS-FM media and plated in a HYPER flask (1720 cm^2^ surface area, Corning). The UCT-MSCs were expanded for 5-7 days, harvested and cryopreserved as P1. Using similar protocol, P1 to P2 expansion was performed, and the cryopreserved P2 UCT-MSC vials were shipped to Georgia Institute of Technology for characterization.

### BM- and AD-MSC Culture

Nine human BM- and three AD-MSCs were purchased from RoosterBio, Inc. (Frederick, MD). Upon purchase BM-MSCs population double level (PDL) ranged from 6.3 – 9.4 and AD-MSCs PDL ranged from 7.81-8.91. These cells were considered as passage 0 (P0). The MSCs were thawed and expanded by protocols provided by RoosterBio, Inc. Cells were cultured in RoosterNourish™ media (High Performance Media Kit, KT-001) in T-225 flasks at an approximate density of 3,333 cells/cm^2^. Cells were grown to 80% confluence (3 – 4 days) and then harvested the following day per manufacturer protocol. Cells were lifted from the flasks by TrypLE Express (Invitrogen) and the enzyme was neutralized with spent media prior to collecting the cells by centrifugation. Cells were frozen in Cryostor CS5 cryopreservation media for future experiments and stored in the vapor phase of liquid nitrogen. Cells were frozen as P1.

### IFN-γ Treatment

P1 BM-MSC donors, BM1 and BM6 were thawed and seeded in T-225 flasks at an approximate density of 3,333 cells/cm^2^. Cells were expanded until they reached ∼90% confluence in RoosterNourish™ media. Cells were then passaged following manufacturing protocols and were plated in 6 – well plates (9 cm^2^) for IFN-γ treatment (approx. 500k cells/well) and allowed for attachment overnight. The media was aspirated and Dulbecco’s Modified Eagle Medium (Thermo Fisher Scientific Inc., Waltham, MA) with 2% fetal bovine serum. The cells were incubated with 50 ng/mL of IFN-γ for 24 hours. After 24-hour IFN-γ incubation, the MSCs were detached with a cell scraper and pelleted for sphingolipid extraction.

### Sphingolipid Extraction

SLs were extracted from MSCs using a modified procedure described elsewhere ^107^. LCBs and CSLs were extracted separately due to their different physical properties. Two aliquots from each MSC sample were taken for LCB analysis and for CSL analysis. An aliquot was also kept for total protein quantification using a BCA protein assay. 1.5 mL of a 2:1 mixture of methanol:methylene chloride was added to each sphingoid base sample and 1.5 mL of a 2:1 mixture of methanol:chloroform was added to each complex sphingolipid sample. Next, 50 pmols of internal standard mixture (Avanti Polar Lipids) was added to each sample. Samples were incubated overnight at 48°C to extract lipids. Next, 150 μL of 1 M KOH in methanol was added to each sample and the samples were incubated at 37°C for 2 hours. This is to cleave the ester bonds of contaminating glycerophospholipids. After incubating, 5 μL of glacial acetic acid was added to all samples to neutralize the KOH. pH was checked using pH strips. 1 mL of chloroform and 2 mL of deionized water were added to all complex sphingolipid samples to induce phase separation. Sphingoid base and complex sphingolipid samples were centrifuged at 1400 xg for 8 minutes to pellet cell debris. For sphingoid base samples, the supernatant was transferred to a new glass tube. For complex sphingolipid samples, the bottom chloroform phase was transferred to a new glass tube. For sphingoid base samples, 0.5 mL of the 2:1 methanol:methylene chloride mixture was added to the cell debris in the original glass tubes. For complex sphingolipid samples, 1 mL of chloroform was added to the cell debris in the original glass tubes. The original tubes were all centrifuged at 1400 xg for 8 minutes. For sphingoid base samples the second supernatant was added to the first supernatant. For complex sphingolipid samples, the second bottom phase was added to the first bottom phase. Remaining cell debris was discarded. Organic solvents were removed by vacuum drying overnight in a Savant SpeedVac (ThermoFisher). The dried SLs were stored in a -20 freezer until analysis.

### Preparation of Samples Sphingolipid Analysis

Dried LCBs samples were resuspended in 300 μL of a 3:2 mixture of mobile phase A1:mobile phase B1 solvent. Mobile phase A1 consisted of 58:41:1 methanol:water:formic acid and 5 mM ammonium formate. Mobile phase B1 consisted of 99:1 methanol:formic acid and 5 mM ammonium formate. Dried complex sphingolipid samples were resuspended in 300 μL of mobile phase A2. Mobile phase A2 consisted of 97:21:1 acetonitrile:methanol:formic acid and 5 mM ammonium formate. Mobile phase B2 consisted of 89:6:5 methanol:water:formic acid and 50 mM triethylammonium acetate. Resuspended samples were centrifuged at 18,000 xg for 10 minutes to remove any remaining cell debris. Total protein amount in each sample was determined for normalization using a Pierce BCA Protein Assay (ThermoFisher Scientific) following the recommended procedure from the manufacturer. The top 200 μL was transferred to a MS autosampler tube.

### Sphingolipid LC-MS/MS Analysis

Sphingolipids were analyzed using Micromass Quattro LC in multiple reaction mode. Fragmented ions were detected with product ion scan. The LCB samples were separated using a 2.1(i.d.) x 150 mm Phenomenex C18 column and a binary solvent system at a flow rate of 300 μL/min. Prior to injection, the column was equilibrated with 100% Mobile phase A1. After injection, the solvent composition was held a 100% A1 for 5 minutes followed by a linear gradient to 100% B1 over 15 minutes. The solvent composition was held at 100% B1 for 5 min, was dropped back to 100% A over 1 minute, and was then held at 100% A for 4 minutes. The CSL samples were separated using a 2.1(i.d) x 150 mm Supelcosil NH2 column and a binary solvent system at a flow rate of 300 μL/min. Prior to injection, the column was equilibrated with 100% mobile phase A2. After injection, the solvent composition was held at 100% A for 5 minutes followed by a linear gradient to 100% B2 for 1 minute. The solvent composition was held at 100% B for 14 minutes, was dropped back to 100% A over 1 minute, and was held at 100% A2 for 9 minutes. Since glucosylceramide and galactosylceramide are not separated by this chromatography method, their concentrations are reported together as hexosylceramide. Internal standards were run to assess retention time drift during MS analysis.

### QUANTIFICATION AND STATISTICAL ANALYSIS

Statistical analysis and False Discovery Rate using the original Benjamini and Hochberg method was conducted in Prism GraphPad 8.0 (GraphPad Software, La Jolla, CA, USA). Spearman Correlation heatmaps were generated using Python matplotlib and Seaborn visualization packages. Comparisons between experimental groups were conducted using Mann-Whitney U-tests and Kruskal-Wallis one-way analysis of variance due to the non-parametric nature of the data. The results are presented as mean ± SEM. JMP Pro 15 was used to for wide linear discriminant analysis (LDA) and principal component analysis (PCA). RStudio (PBC, Boston, MA) was used to generate graphical illustrations of PCA variable plot. Montgomery

