## Supplementary Figure 1 for "Characterizing Human Mesenchymal Stromal Cells Immune Modulatory Potency Using Targeted Lipidomic Profiling of Sphingolipids" (from another version of this preprint: version 2, 2021-06-07)

**SUPPLEMENTARY INFORMATION**

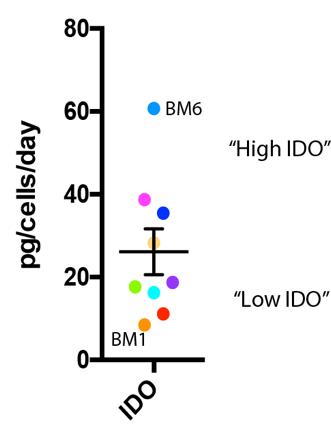

Supplementary Figure 1. Principal component analysis plot of bone-marrow derived MSC’s indoleamine-2,3-dioxygenase activity. IDO values above the mean were classified as “high IDO”. IDO values below the mean were classified as “low IDO” (N = 9).
